# Modes of inhibition used by phage anti-CRISPRs to evade type I-C Cascade

**DOI:** 10.1101/2022.06.15.496202

**Authors:** Roisin E. O’Brien, Jack P.K. Bravo, Delisa Ramos, Grace N. Hibshman, Jacquelyn T. Wright, David W. Taylor

**Affiliations:** Interdisciplinary Life Sciences Graduate Programs, University of Texas at Austin, Austin, TX, 78712; Department of Molecular Biosciences, University of Texas at Austin, Austin, TX, 78712; Center for Systems and Synthetic Biology, University of Texas at Austin, Austin, TX, 78712; LIVESTRONG Cancer Institutes, Dell Medical School, University of Texas at Austin, Austin, TX, 78712

## Abstract

Cascades are RNA-guided multi-subunit CRISPR-Cas surveillances complexes that target foreign nucleic acids for destruction. Here, we present a 2.9-Å resolution cryo-electron (cryo-EM) structure of the *D. vulgaris* type I-C Cascade bound to a double-stranded (ds)DNA target. Our data shows how the 5’-TTC-3’ protospacer adjacent motif (PAM) sequence is recognized, and provides a unique mechanism through which the displaced, single-stranded non-target strand (NTS) is stabilized via stacking interactions with protein subunits in order to favor R-loop formation and prevent dsDNA re-annealing. Additionally, we provide structural insights into how diverse anti-CRISPR (Acr) proteins utilize distinct strategies to achieve a shared mechanism of type I-C Cascade inhibition by blocking initial DNA binding. These observations provide a structural basis for directional R-loop formation and reveal how divergent Acr proteins have converged upon common molecular mechanisms to efficiently shut down CRISPR immunity.

## Introduction

CRISPR (clustered, regularly, interspaced, short palindromic repeats) and CRISPR-associated (Cas) genes together form a prokaryotic adaptive immune response against foreign genetic elements, such as plasmids and phages^1, 2^. CRISPR-Cas immunity is established through three major stages: adaptation, maturation, and interference^3, 4^. Immunological memory is first acquired through the incorporation of short genetic fragments from invading phage or plasmids into the bacterial genome at the CRISPR loci^3, 4^. These short fragments are then transcribed into pre-CRISPR RNA (crRNA) and processed into mature crRNA transcripts^5^. Cas proteins assemble around the mature crRNA to form either multi-subunit or single subunit crRNA-guided surveillance complexes, utilizing the crRNA as a guide for target recognition^6^. Once base-complementarity between the crRNA and the target has been established and a protospacer adjacent motif (PAM) has been identified, CRISPR effector complexes cleave and degrade these nucleic acid targets^7^. Despite this elegant defense system, phages evade CRISPR-Cas complexes by counter attacking with small inhibitory proteins known as anti-CRISPRs (Acrs)^8, 9^. The interplay between anti-CRISPRS and CRISPR-Cas systems is one component of the molecular arms race between phage and bacteria^10^.

CRISPR-Cas systems are highly diverse and can be divided into two major classes^11, 12^. Class I use a multi-subunit crRNA-guided surveillance complexes, while Class II rely on a single effector nuclease^13, 14^. While the simplicity and programmability of Class II nucleases has generated major attraction for genome engineering applications, these systems are present in less than 10% of bacteria and archaea^12^. Strikingly, Class I systems account for almost 90% of CRISPR-Cas systems observed in nature. This class can be further divided into types I, III, and IV^12^. Among all CRISPR-Cas systems, type I is the most prevalent. Type I systems are characterized by the presence of a trans-acting helicase-nuclease, Cas3, and divided into multiple subtypes (A-G)^12, 14^.

Interestingly, the type I-C CRISPR-Cas system contains only three unique Cas proteins in its operon: Cas5c, Cas7, and Cas8c^12^. Accordingly, it is a minimal Cascade. The type I-C Cascade uses Cas5c for processing the crRNA instead of a separate Cas6^12, 15–17^ and does not include a small subunit (Cas11c) within its operon^12^. Early structural studies hypothesized that the large subunit, Cas8c, was a fusion of the larger and smaller subunits found in the type I-E Cascade^16^. However, recent studies revealed that the *D. Vulgaris* large subunit Cas8c includes an internal ribosome binding site at the C-terminus, which encodes a separate small subunit Cas11c^18^. We previously confirmed that this non-canonical Cas11c is identical to the C-terminal domain of Cas8c in both sequence and structure, adopting a helical bundle topology typical of other small subunits^17^. The discovery and inclusion of the Cas11c small subunit has been a crucial component for effectively utilizing type I-C systems for genome engineering applications^19^.

While previous structures and biochemical analysis of the type I-C Cascade have provided insights into complex assembly^16, 17^, little information exists about how type I-C Cascade recognizes targets. The growing use of type I-C Cascade for genome editing applications underlines the importance for a mechanistic understanding of PAM recognition and dsDNA unwinding and stabilization during R-loop formation^19, 20^. Furthermore, modulating type I-C *in vivo* functionality though the incorporation of anti-CRISPR (Acr) inhibitory proteins is also of interest^19, 21^. The lack of sequence similarity and structural motifs among anti-CRISPR proteins has hindered their identification, including those targeting the I-C effector complex^9^. However, a novel method for discovering anti-CRISPRs recently revealed multiple type I-C anti-CRISPR proteins that inactivated the I-C effector complex *in vivo,* most notably AcrIC4^22^. Surprisingly, a previously characterized type I-F inhibitory protein, AcrIF2, demonstrated *in-vivo* inhibition of the type I-C Cascade as well^22, 23^. We were interested in understanding the mechanistic details that fueled the dual *in-vivo* inhibitory activity of AcrIF2 and compare it to a novel I-C specific anti-CRISPR, AcrIC4.

Here, we use cryo-electron microscopy (cryo-EM) to determine a 2.9-Å structure of the type I-C Cascade bound to a double-stranded (dsDNA) target. This structure reveals dramatic conformational changes that are crucial for R-loop formation and novel mechanisms of non-target strand stabilization and PAM recognition. Additionally, we determined structures of the type I-C Cascade bound to AcrIF2 and AcrIC4 at 3.0-Å and 3.1-Å resolution, respectively. These structures unveil the different strategies AcrIF2 and AcrIC4 use to achieve inhibition of type I-C Cascade PAM recognition and subsequent dsDNA binding. Collectively, this study expands our mechanistic understanding of the type I-C effector complex and provides insights into the modes of inhibition Acrs use to help phage evade type I CRISPR bacterial systems.

## Results

### DNA Binding Induces Conformational Changes in the Type I-C Cascade

To understand the mechanism of PAM recognition and R-loop formation by type I-C Cascade, we determined a 2.9-Å cryo-EM structure of this effector from *D. vulgaris* bound to a 75-bp double stranded DNA (dsDNA) target containing a complimentary protospacer and 5’-TTC PAM (Figure 1a-c and S1-2). To assist with modeling, we improved our previously-determined^17^ apo Cascade cryo-EM structure from 3.1- to 2.7-Å from this same dataset, enabling us to unambiguously model several regions of the complex that were poorly-resolved in our previous structure. Due to molecular motion, residues 290-348 within the Cas8c subunit are missing in both the apo and DNA-bound structures. In the DNA-bound structure, a combination of density subtraction, focused 3D classification and local refinement significantly improved the resolution of the flexible Cas8c N-terminus, and AlphaFold2 was used to further assist with modeling of a full R-loop structure (Figure1e left and S1-2)^24, 25^. Incidentally, we observed a subset of particles within our cryo-EM dataset that lacked the non-target strand (NTS), but formed a hybrid between the crRNA spacer and the DNA target strand (TS), which resulted in a structure at 2.9-Å resolution. Interestingly, the Cas8c N-terminus was highly flexible and absent in our structure, akin to the apo Cascade structure.

**Figure 1.**
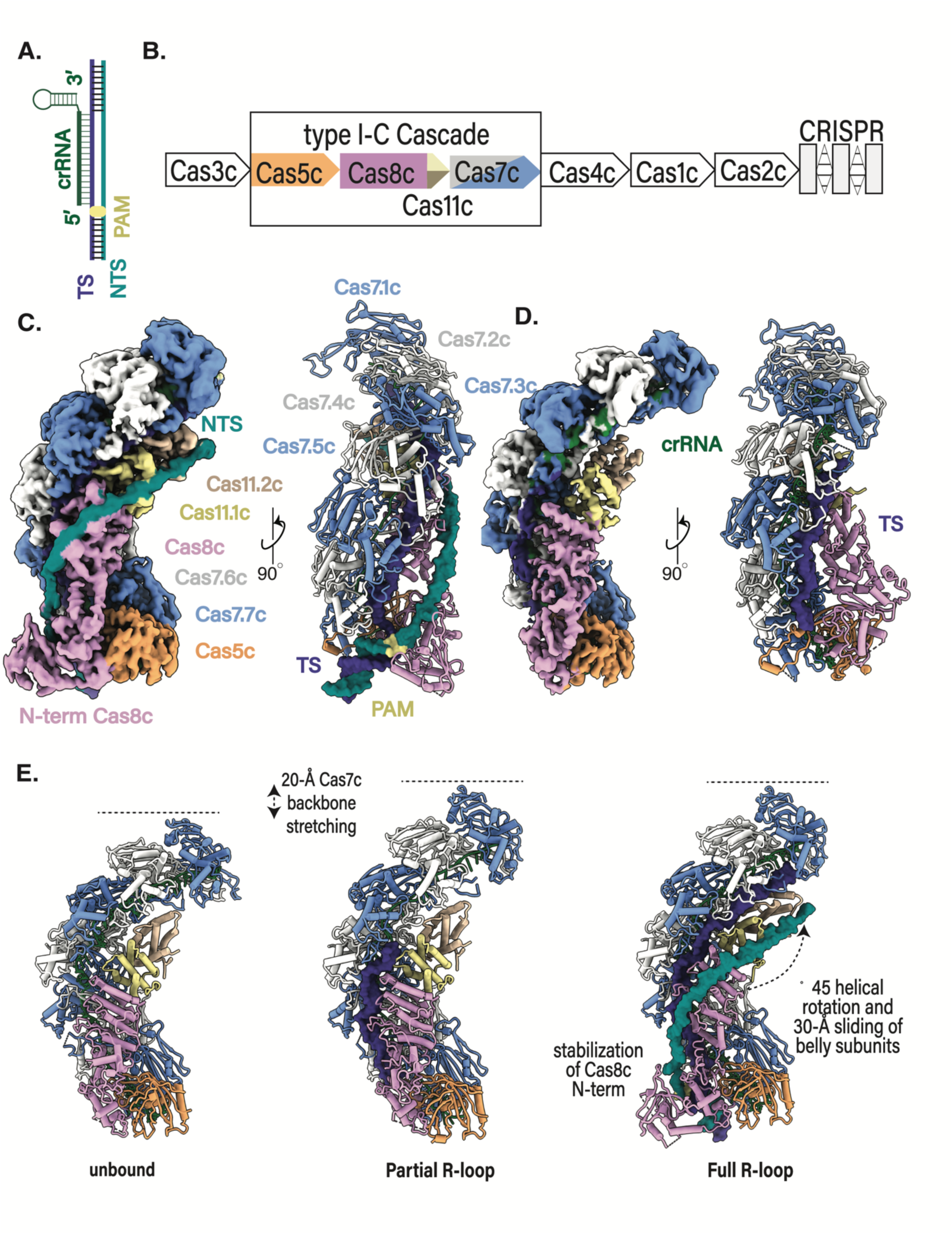
Structural Confirmations of the *D. vulgaris* type I-C Cascade. **a,** R-loop schematic of dsDNA target. **b,** Gene organization of the D. *vulgaris* type I-C operon. **c,** 2.9-Å cryo-electron (cryo-EM) structure and atomic model of the type I-C Cascade bound to a double-stranded (ds)DNA target **d,** 2.80-Å cryo-electron (cryo-EM) structure and atomic model of the type I-C Cascade bound to a single-stranded (ds)DNA target**. e,** Conformational changes of type I-C Cascade induced upon ssDNA and dsDNA binding compared to improved 2.7-Å apo-model.

The overall architecture of type I-C Cascade with or without either target DNA consistently resembles a caterpillar, comprised of five unique Cas proteins in Cas7c_7_ Cas5c_1_Cas8c_1_Cas11c_2_ stoichiometry and a single 35-nt crRNA^16, 17^ (Figure 1c-e). The complex first assembles with the crRNA-processing subunit, Cas5c, sitting at the base of the complex and cradling the 5’-end of the crRNA handle^16, 17^. Seven Cas7s stack on top of the Cas5c, oligomerizing along the crRNA in a helical filament until they cap the 3’-end of the crRNA^16, 17^. Finally, the large Cas8c subunit and two non-canonical Cas11 small subunits nestle inside the belly of the complex, completing the arrangement of subunits of this effector^17, 18^. In both the ssDNA structure and dsDNA structures, the entire target strand (TS) runs parallel to the complimentary crRNA, wedged inside the complex (Figure 1c-d). When bound to dsDNA, the non-target strand (NTS) is guided along the outside of the complex by the Cas8c and Cas11c belly subunits (Figure 1c).

While still maintaining the same overall caterpillar architecture, the type I-C Cascade undergoes multiple conformational rearrangements to achieve full R-loop propagation (Figure 1e). The Cas7-crRNA backbone becomes extended in the ssDNA and dsDNA bound structures, stretching by ∼20Å to accommodate base pairing of the crRNA to the TS protospacer (Figure 1e) ^17^. Additionally, the N-terminus of Cas8c (Cas8c N-term) is stabilized upon dsDNA binding, interacting with the NTS at the base of the complex exclusively when bound to dsDNA (Figure 1c,e)^17^. However, the most dramatic conformational change involves a largely uniform rigid-body rearrangement of the belly subunits. dsDNA binding causes the Cas8c C-terminal domain and the two Cas11c subunits to shift upwards by 20-Å and rotate by ∼90°. Since these conformational changes are exclusive to the dsDNA-bound structure, it is likely that they occur by allosteric signaling within Cascade upon recognition of a suitable PAM and accompany R-loop propagation. (Figure 1e).

### Cas8c demonstrates novel mechanisms for NTS stabilization

Previous type I Cascade structures captured with complete R-loops had limited resolution of the NTS, preventing direct visualization of how the NTS is stabilized during R-loop propagation. This mechanism is critical for kinetically partitioning the forward and backward reactions, preventing dsDNA reannealing and favoring R-loop completion. Since charge-swap mutations introduced at this site resulted in a substantial DNA-binding defect for type I-E and I-F Cascades^26, 27^, it has been proposed that the NTS is stabilized predominantly through non-specific electrostatic interactions with the surface of the small and large subunits^26, 27^. Our dsDNA-bound Cascade structure exhibited excellently resolved cryo-EM density for the displaced NTS, enabling us to confidently model 41nts of the NTS and providing a structural snapshot of the interactions between NTS and Cascade (Figure 2). As previously hypothesized, positively charged residues (R205, R408, K459, K607, K608) on the surface of the C-term of Cas8c participate in non-specific electrostatic interactions with the negative NTS backbone^26, 27^ (Figure 2a). Because Cas11c is identical in sequence and structure to the C-terminus of Cas8c, the analogous positively charged residues in the Cas11 subunits (K118) interact with the phosphate backbone until the NTS reanneals with the TS at the top of the complex (Figure 2b). Interestingly, the only highly conserved residues across type I-C subtypes are K607 and K608 (Data S1). Due to the lack of sequence-specific interactions with the NTS, however, it is likely that the surface of Cas8c-Cterm and Cas11 containing a positively charged channel accommodates the NTS, removing evolutionary selection and explaining the lack of residue conservation for this region.

**Figure 2.**
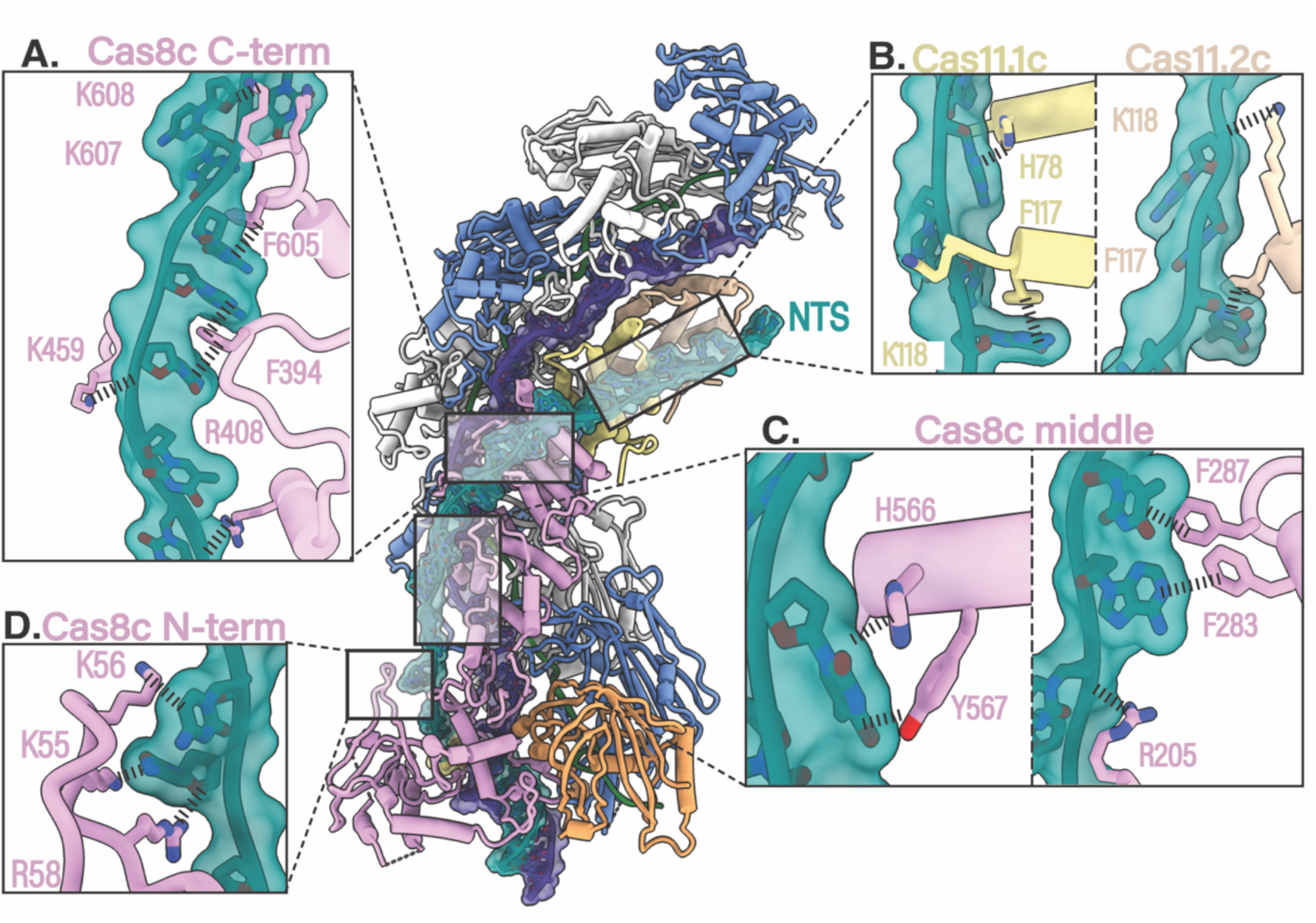
Cas8c demonstrates novel mechanisms for NTS stabilization. **a-c,** Various residues in Cas8c and Cas11 small subunits are involved in NTS stabilization. Positively charged and aromatic residues across surface of the Cas8c and Cas11 subunits form non-specific interactions with the NTS backbone and bases, respectively. **d,** Cas8c N-term acts a dsDNA “vice” that folds in around the dsDNA helix and facilities initial melting of the duplex using highly conserved, positively charged residues to hold the NTS and prevent reannealing with the TS

Intriguingly, we identified a novel mechanism of NTS stabilization whereby aromatic residues on the surface of the Cas8c C-terminus (F283, F287, F394, H566, Y567, F605) participate in stacking interactions with NTS bases. This may have not been obvious in previous Cascade structures due to limited resolution in this area^26–28^. Again, because of the non-canonical nature of the type I-C small subunits, the analogous residues in Cas11 (H78 and F117) continue to form the same base-stacking interactions towards the PAM-distal end of the NTS (Figure 2b). Residues F283, F287, F394, F605 were highly conserved among type I-C Cascades, where positions 566 and 567 displayed conservation of an aromatic residue (Tyr, Phe, or His) (Data S1). Strikingly, we identified multiple aromatic residues similarly positioned along the putative NTS path of type I-E Cascade^27^ (Figure S3), suggesting that this mechanism of NTS stabilization may occur within multiple type I systems.

At the PAM-proximal end, the NTS is first pulled apart from the duplex through a series of positively charged residues emerging from a loop protruding from the Cas8c N-terminus (Figure 2d). Similar to type I-F Cascade, the Cas8c N-term acts a dsDNA “vice” that folds in around the dsDNA helix and triggers initial melting of the duplex^26^ (Figure 2d). Highly conserved, positively charged residues (K55, K56, R58) within the Cas8c “vice” make sequence-independent contacts with the negatively charged NTS backbone (Figure 2d, Data S1). Since the N-terminus of the Cas8c could not be resolved in the apo or ssDNA structure, these charge-charge contacts between the Cas8c N-terminus and the NTS play a role in stabilizing the NTS during R-loop formation^17^.

### Cas8c N-term is responsible for PAM recognition

Type I Cascades recognize target sequences through both non-sequence specific contacts and sequence-specific contacts with a protospacer adjacent motif (PAM)^26–28^. The PAM is a short nucleotide motif upstream of the target sequence that facilitates distinction of self from non-self DNA^29^ (Figure 1a). In the type I-C Cascade, the 5’-TTC PAM motif is recognized from the minor grove by the N-term of the Cas8c large subunit (Figure 3a). The N-term of Cas8c clamps around the dsDNA helix and positions four loops to facilitate PAM recognition and strand separation (Figure 3b-c). Asparagine 72 (N72) makes the first contact with the PAM, protruding from a glycine loop that wedges between the A_T-3_:T_NT-3_ base pairs (Figure 3c,d). N72 is within hydrogen bonding distance to both the TS A_T-3_ and the NTS T_NT-3_ (Figure 3b). The N72 creates dual contacts with both strands and supports strict tolerance for only pyrimidines at PAM NT_-3_ position for some type I-C Cascades, yet is intriguingly not a highly conserved residue across type I-C Cascades (Data S1). While a glycine loop is a PAM recognition feature present in type I-E Cascades, the use of asparagine for PAM recognition bares more of a resemblance to type I-F Cascade^26–28^. To verify the role of N72 in PAM recognition, we introduced an alanine mutation at this position and performed electrophoretic mobility shift assays (EMSA) with both WT and N72A Cas8c containing Cascades (Figure 3f-g). Compared to WT, N72A had significantly decreased DNA binding affinity, supporting its important contribution to PAM recognition (Figure 3g).

**Figure 3.**
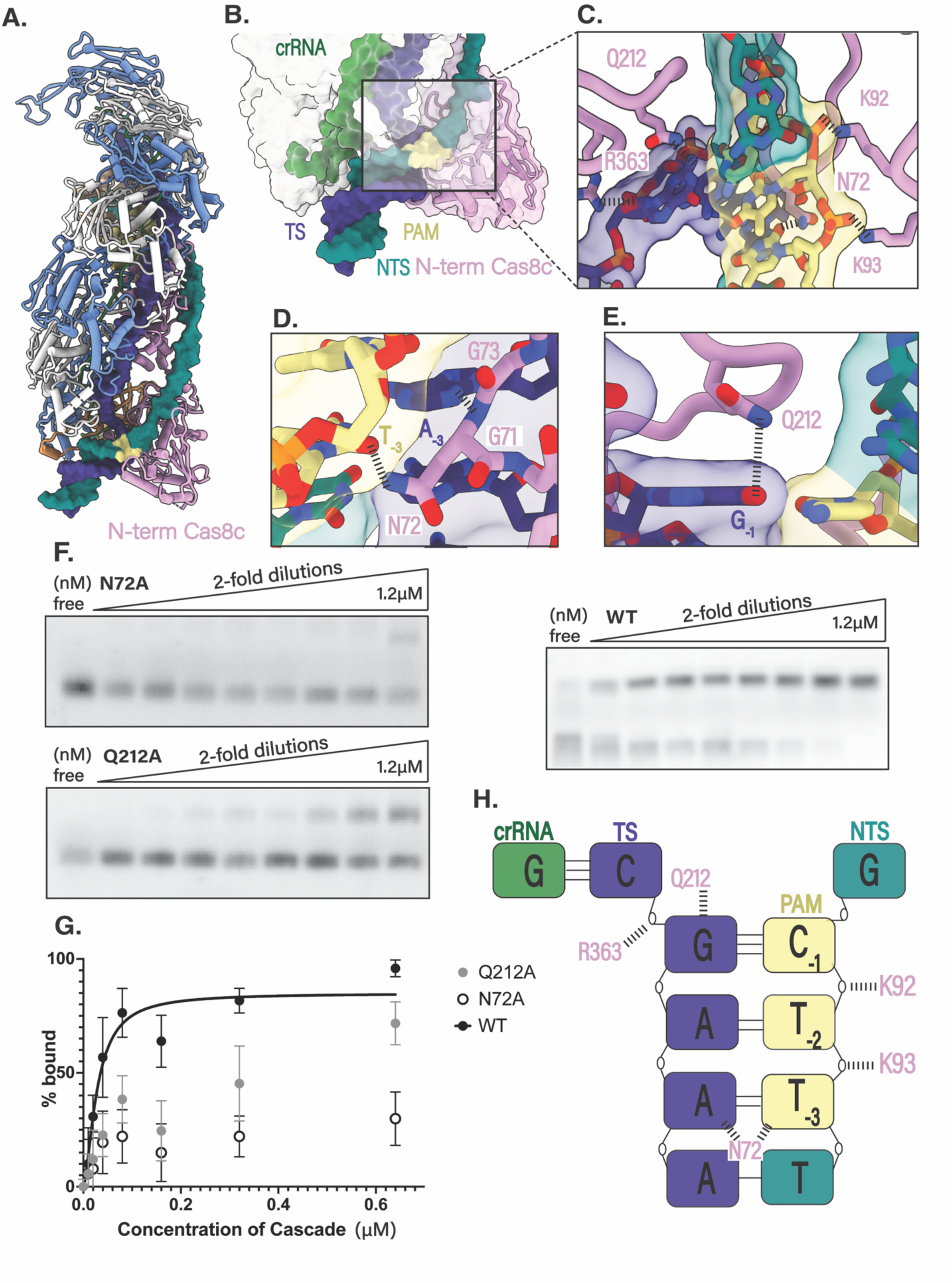
Cas8c N-term presents a more minimal PAM recognition scheme. **a,** 5’-TTC PAM motif is recognized from the minor grove by the N-term of the Cas8c large subunit **b,** The N-term of Cas8c clamps around the dsDNA helix and **c,** positions four loops to facilitate PAM recognition and strand separation. **d,** Asparagine 72 (N72) makes the first contact with the PAM, protruding from a glycine loop that wedges **e,** dsDNA splitting is enabled by a glutamine wedge that stacks above the PAM, and intercalates between the two DNA strands **f,** Electrophoretic mobility shift assays (EMSA) measuring binding of increasing concentrations of Cascade to a dsDNA target with a 5’-TTC PAM. Experiments were done in triplicate and representative results are shown. **g,** Quantification of gel shift data. Each point is the average of at least three independent replicates. Binding curves for mutants are omitted due to poor fit. WT Cascade Kd is 28nM. **h,** Summarization of the five residues involved in the PAM recognition mechanism for the type I-C Cascade. Two residues (Q212A and N72A) make specific contact with the PAM bases and three residues (K92, K93, R363) stabilize the PAM phosphate backbone

Like type I-E Cascades, dsDNA splitting is enabled by a glutamine wedge that stacks above the PAM, and intercalates between the two DNA strands^27, 28^ (Figure 3e). Q212 hydrogen bonds with G_T-1,_ sterically displacing the first two nucleotides of the protospacer and forcing them to rotate outwards. Given its strategic location, it is not surprising that Q212 is highly conserved across type I-C Cascades (Data S1). To test the requirement of this wedge for target binding, we introduced an alanine mutation at the Cas8c Q212 position and measured its impact using EMSAs. The Q212A effector mutant resulted in a decrease in DNA binding affinity compared to the WT (Figure 3f-g). The final two loops involved in PAM recognition contain three positively charged residues (K92-92, R363) that non-specifically stabilize the PAM duplex phosphate-backbone (Figure 3h).

### AcrIF2 blocks PAM recognition and domain rearrangements required for activation

Phages can utilize anti-CRISPR proteins (Acrs) to deactivate CRISPR-Cas effectors and escape detection^8, 9^. AcrIF2 has demonstrated high levels of CRISPRi suppression *in vivo* across multiple type I subtypes^22^. To determine the mechanism of AcrIF2 inhibition towards the type I-C Cascade, we determined a 3.0-Å cryo-EM structure of AcrIF2 bound to the type I-C Cascade (Figure 4a and S1-2). Binding of the AcrIF2 does not generate any conformational displacement of the type I-C Cascade subunits when compared to the apo model (Figure 4a). However, density for the Cas8c N-term was present, as observed only in the dsDNA bound cryo-EM map (Figure 1c and 4a inset). A single AcrIF2 sits at the hinge of Cas8c, between the N-terminus and the rest of the subunit (Figure 4a-b). It interacts with a positively charged interface of Cas8c through a large network of electrostatic, making no contacts to any of the other subunits (Figure 4a-b). Compared to a previous cryo-EM structures of AcrIF2 bound to a type I-F Cascade, AcrIF2 uses the same acidic interface to bind both Cas8f and Cas8c^23^ (Figure 4c, S4). However, AcrIF2 is oriented parallel rather than perpendicular to Cas8c (i.e. is rotated by 90°), which provides additional contacts with Cas8c than Cas8f^23^ (S4). Structural superposition of the type I-C dsDNA bound model shows severe clashing between PAM and AcrIF2 (Figure 4d). AcrIF2 sits right at the PAM site, both sterically blocking the PAM sequence, but also interacting with a crucial residue involved in PAM recognition (Q212) (Figure 4d). Not only does AcrIF2 crowd the PAM site, but it is large enough to also prevent the Cas8c N-term from clamping around the dsDNA target (Figure 4d-e). As we previously demonstrated, the N-term is essential for dsDNA binding, involved in both PAM-recognition and NTS stabilization (Figure 2g). Thus, AcrIF2 engages in a two-pronged attack to prevent dsDNA binding of the type I-C Cascade - it wedges within the PAM-binding site of Cas8c, but it additionally jams the vice-like C-terminus in a non-productive configuration, further abrogating any possibility of PAM-recognition (Figure 4d-e). To test the inhibition of AcrIF2, we performed *in vivo* cleavage assays measuring type I-C Cascade-Cas3 cleavage activity in the presence of IF2 (Figure 4f). Consistent with our structural hypothesis and previously corroborating *in vivo* data, AcrIF2 effectively blocks target dsDNA binding to the type I-C Cascade by at least two fold^21^ (Figure 4f).

**Figure 4.**
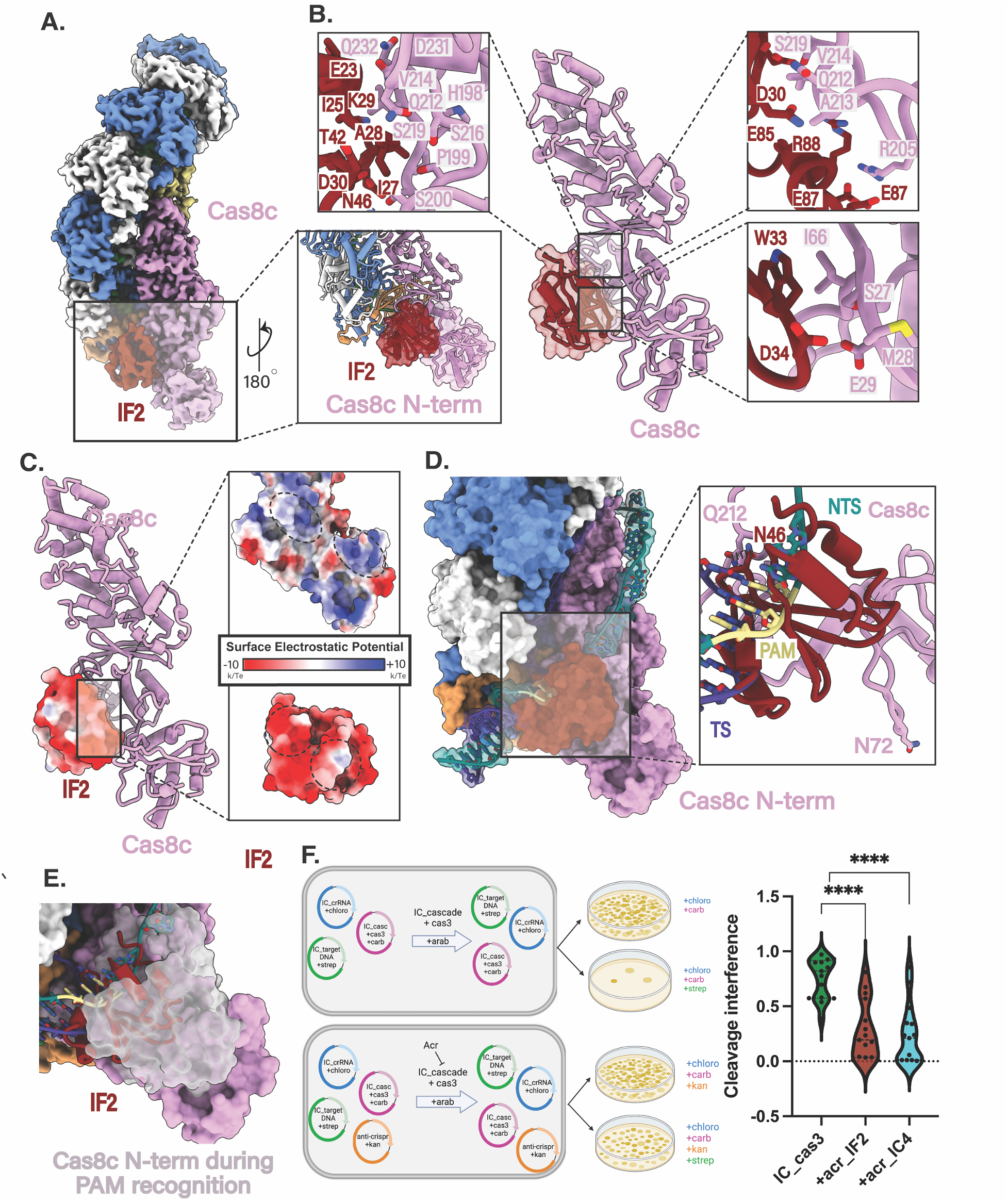
AcrIF2 blocks both PAM recognition and domain rearrangements required for activation. **a,** 3.0-Å cryo-electron structure of the type I-C Cascade bound to AcrIF2 and atomic model demonstrating the presence of Cas8c N-term (inset) **b,** AcrIF2 exclusively interacts with Cas8c through a large network of non-specific interactions **c,** AcrIF2 is entirely negative and and sits within two positively-charge surfaces between Cas8c N-term and the rest of the Cas8c subunit **d,** Structural superposition of the type I-C dsDNA bound model shows severe clashing between PAM residues and AcrIF2. AcrIF2 additionally hinders the rearrangement of Cas8c N-term containing important residues involved in PAM recognition (N72). **e,** AcrIF2 prevents the Cas8c N-term dsDNA “vice” (grey) from folding in around the dsDNA helix and facilitating initial melting of the duplex (pink) **f**, type I-C Cascade-Cas3 *in vivo* interference assays demonstrate AcrsIF2 and AcrIC4 effectively blocks dsDNA binding by more than 2-fold

### AcrIC4 causes steric hindrance at the PAM site

Unlike AcrIF2, AcrIC4 has demonstrated the highest levels of CRISPRi suppression *in vivo,* exclusively targeting the type I-C subtype^22^. To understand the mechanism of this type I-C Cascade-specific Acr, we solved a 3.1-Å cryo-EM structure of the type I-C Cascade bound to AcrIC4 (Figure 5a and S1-2), and used AlphaFold2 to assist with structural modeling. A single AcrIC4 lies at the base of the type I-C Cascade, wedged in between Cas7.6c and Cas8c (Figure 5a-b). Binding of IC4 generates no conformational change of the type I-C Cascade subunits compared to the apo model (Figure 5a). A negatively charged interface on IC4 interacts with a positively charged surface on both the Cas7.6c and Cas8c subunits through primarily hydrophobic and charge-charge contacts (Fig 5b and 5c). Structural superposition of the dsDNA-bound model shows severe clashing between the PAM and AcrIC4 (Figure 5d). AcrIC4 not only sterically blocks dsDNA binding at the PAM, but it also lies within hydrogen bonding distance of a crucial PAM-recognizing residue of Cas8c (Q212) (Figure 5d inset). Based on the lack of density for the Cas8c N-term, AcrIC4 does not engage with the N-term of Cas8c in a similar manner to AcrIF2 (Figure 4a and 5a). However, even with Cas8c N-terms ability to engage with the PAM site, the structural interference AcrIC4 poses for PAM recognition is likely insurmountable for achieving dsDNA binding (Figure 5d inset). Interestingly, although AcrIF2 and AcrIC4 utilize similar strategies to inhibit type I-C Cascade (i.e. blocking PAM binding), both proteins do so by targeting slightly different (but partially overlapping) surfaces of the complex. This suggests that while PAM blocking is a widespread anti-CRISPR strategy, Acrs use diverse strategies to target hotspots within Cascade (Figure 5e). To test the inhibition of AcrIC4, we performed *in vivo* cleavage assays measuring type I-C Cascade-Cas3 cleavage efficiency in the presence of AcrIC4 (Figure 4f). Consistent with our structural hypothesis and previous corroborating *in vivo* data, AcrIC4 effectively blocks target dsDNA binding to the type I-C Cascade by at least two-fold.^21^

**Figure 5.**
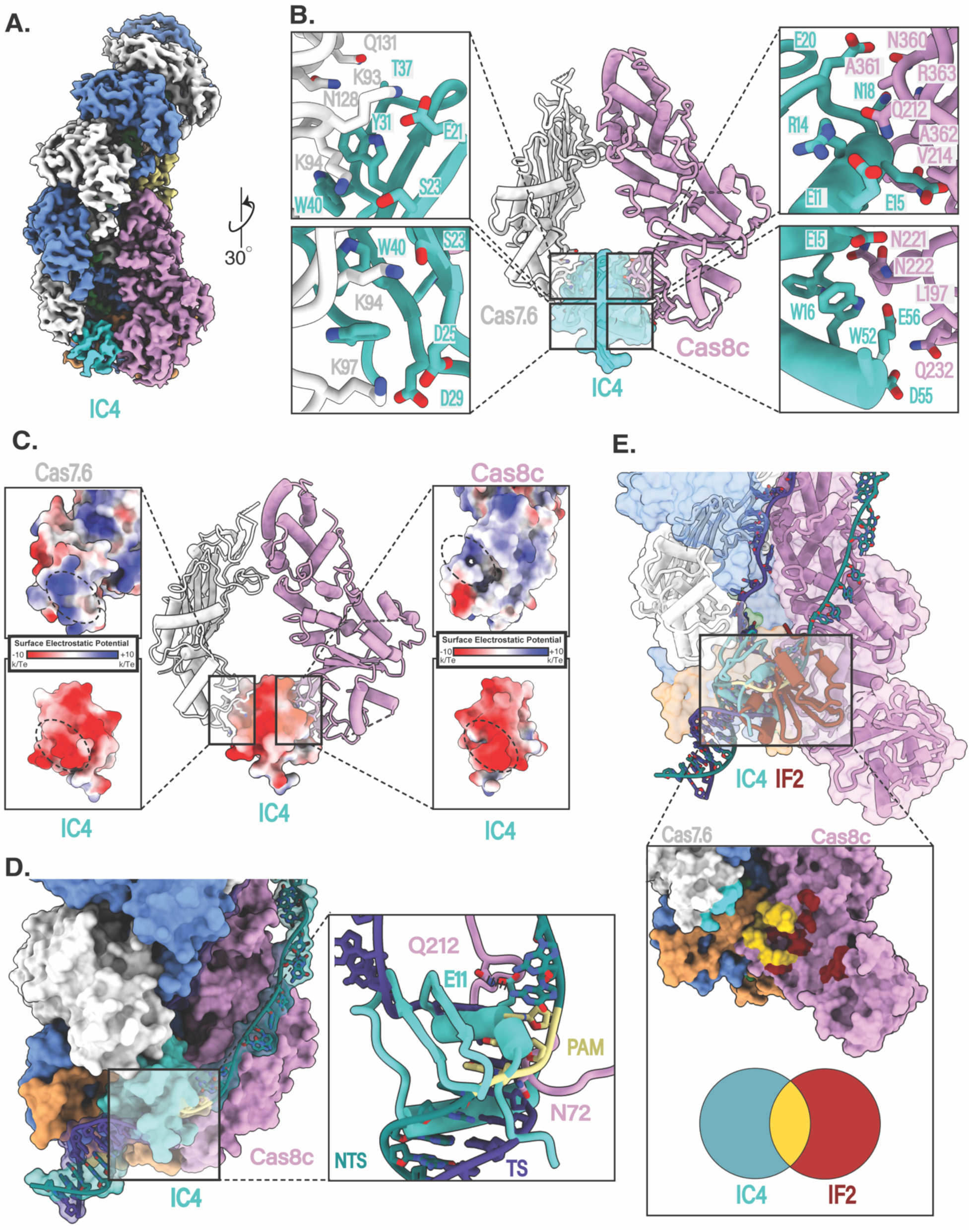
AcrIC4 causes steric hindrance at the PAM site. **a,** 3.1-Å cryo-electron structure of the type I-C Cascade bound to AcrIC4 **b,** AcrIC4 interacts with both Cas8c and Cas7c through a large network of non-specific interactions **c,** AcrIC4 is entirely negatively charged and wedged between positively-charged Cas8c and Cas7.6 surfaces **d,** Structural superposition of the type I-C dsDNA bound model shows severe clashing between PAM residues and AcrIC4. **e,** Overlay of the IF2 and IC4 binding sites demonstrate an overlapping interface at the PAM site.

## Discussion

Here, we describe the mechanisms by which the type I-C Cascade is activated for dsDNA targeting and how two anti-CRISPR proteins can deactivate it. Three high-resolution cryo-EM structures of Cascade at different stages of activation reveal critical details about how target DNA binding triggers major conformational rearrangements for R-loop formation. While DNA target binding induces complex elongation, recognition of a cognate PAM sequence and R-loop propagation are essential to trigger conformational changes in the C-terminus of Cas8c and Cas11c subunits, locking the complex into a complete R-loop. Similar phenomena have been observed for the type I-F Cascade^26^. Thus, our structures highlight the importance of duplex vs ssDNA binding for type I systems in triggering R-loop formation^26–28^ and establishing conformational control mechanisms to ensure correct target recognition for Cas3 recruitment and activation^30, 31^.

Despite the importance of the NTS in creating a stable R-loop, a thorough understanding of the structural mechanisms behind NTS stabilization were previously hindered by limited resolution in this region in other structures^26–28^. Here, we present 41-nt of the NTS, which enabled full visualization of interactions between the NTS and the Cas8c and Cas11c belly subunits (Figure 2). For the type I-C Cascade, NTS stabilization is built upon a multi-layer network of support as a structural foundation. As prior type I Cascade structures have alluded to, the first layer is comprised of positively-charged residues that non-specifically contact the NTS phosphodiester backbone^26–28^. A second layer engages aromatic residues to ‘pinch’ the bases of the NTS, reinforcing rigidity by fastening the nucleotides into place. Interestingly, the type I-E Cascade contains similar aromatic residues located along the NTS pathway that would be ideal for providing similar contacts^27^. Thus, the combination of charge-charge and base-stacking interactions underpins the importance of securing the NTS during R-loop formation across type I CRISPR-Cas surveillance complexes.

Our structural and biochemical data provides valuable insights into PAM recognition. While clear characteristics with other Cascades are evident, type I-C demonstrates a more minimal PAM recognition scheme compared to its other type I counterparts. It requires only two residues (N72 and Q212) to interact with PAM bases and is a unique mechanism for type I Cascades. Interestingly, type I-C systems have consistently demonstrated a strong preference for the 5’-TTC-3’ PAM *in vivo*^32^. Yet, for such a strict tolerance, fewer interactions are required to recognize the PAM by the Cas8c subunit, compared to those of other Cascades. Recent structural studies on the type I-C acquisition complex showed that Cas4 incorporates several PAM-recognition residues, which results in a particularly stringent PAM-recognition mechanism^33^. Thus, we hypothesize that the strong preference of a type I-C 5’-TTC PAM is likely perpetuated during acquisition, rather than the interference stage of CRISPR-Cas immunity. However, future studies will be necessary to fully elucidate these mechanisms. It is tempting to hypothesize that this minimal PAM-recognition mechanism could be used for engineering a minimal type I Cascade with a more promiscuous PAM tolerance.

Finally, we present the first structures of the type I-C Cascade bound to AcrIF2 and AcrIC4. Through our structural analysis, we were able to show the different strategies Acrs use to achieve the same goal of blocking PAM binding (Figure 3 and 4). AcrIC4 interacts with both Cas7c and Cas8c subunit and bocks dsDNA binding by acting as a negatively-charged structural blockade at the PAM site (Figure 5). In contrast, AcrIF2 wedges into the positively charged hinge of the Cas8c subunit, creating bipartite steric hindrance of both the PAM site and the N-term of Cas8c (Figure 4). Previous structural analysis of AcrIF2 proposed that this anti-CRISPR was representative of a “DNA mimic,” however, we believe that the mechanism of AcrIF2 is more complex in type I-C Cascade^23^. The entirety of the AcrIF2 is negatively charged, which enables it to wedge into the positively charged hinges of Cas8 subunits across both type I-C and type I-F subtypes. And because of AcrIF2’s relatively large size compared to AcrIC4 (10kDa to 6Kda, respectively), it creates a second level of inhibition by holding the Cas8c N-term away from stable PAM clamping and R-loop propagation. Collectively, these structures outline two strategies used among anti-CRISPRS to achieve the common mechanism of blocking PAM binding. These structures could provide the basis for improving type I-C Cascade as a genome engineering tool and understanding how to best regulate them using Acrs for these applications.

## Methods

### Generation of AcrIC4 Plasmid

The AcrIC4 protein sequence (WP_153575361.1)^22^ was codon optimized and cloned into the pET His6 SUMO vector (#48313) using ligation independent using the primers listed in Table 3. Fusion tags are added to the 5’ end of each primer. All associated plasmids and vectors can be found in Tables 2 and 3, respectively.

### Oligonucleotide Preparation

DNA oligonucleotides used in cleavage assays, gel shifts, and electron microscopy were purchased from Integrated DNA Technologies. The dsDNA duplex was formed by mixing equimolar TS and NTS (Table 3) in 40 mM Tris (pH 8.0) and 38 mM MgCl2 heating at 95°C for 2 min, and slow cooling at room temperature for at least 10 min^16^.

### Protein Purification

The WT and PAM mutants of D*. vulgaris* type I-C Cascade (addgene plasmid #81185) were co-expressed with crRNA (addgene plasmid #81186) in NiCo21(DE3) *E. coli* cells^16^. AcrIF2 (addgene plasmid #89234)^23^ and AcrIC4 were also co-expressed in Nico21(DE3) *E. Coli* cells and follow the same following purification protocol as I-C Cascade. Cells were grown at 37°C to an OD600 of 0.6-0.8 and induced by the addition of 0.5 mM isopropyl-β-D-thiogalactopyranoside (IPTG). After overnight growth at 18°C, the cells were harvested and lysed by sonication in a buffer containing 50 mM HEPES–NaOH (pH 7.5), 500 mM KCl, 5% (v/v) glycerol, 1 mM tris(2-carboxyethyl)phosphine (TCEP), 0.01% Triton X-100, 0.5 mM PMSF, and complete Roche mini protease inhibitor tablets. The lysate was centrifuged at 27,000 × g and applied to a HisTrap Ni-NTA affinity column, pre-equilibrated in lysate 50 mM HEPES–NaOH (pH 7.5), 500 mM KCl, 5% (v/v) glycerol, 1 mM tris(2-carboxyethyl)phosphine (TCEP). The protein-bound resin was washed with buffer containing 50 mM HEPES–NaOH (pH 7.5), 150 mM KCl, 5% (v/v) glycerol, and 1 mM TCEP and a second buffer containing 20 mM imidazole, 50 mM HEPES–NaOH (pH 7.5), 150 mM KCl, 5% (v/v) glycerol, and 1 mM TCEP. Protein was eluted with 50 mM HEPES– NaOH (pH 7.5), 150 mM KCl, 5% (v/v) glycerol, 1 mM TCEP, and an imidazole gradient up to 500mM. Approximately 1 mg of TEV protease was added per 25 mg of protein and the protein-TEV mixture was dialyzed at 4°C overnight against size-exclusion buffer. The protein was then concentrated to approximately 20mg/ml and run over a Superdex 200 Increase 10/300 GL size-exclusion column in a buffer containing 50 mM HEPES–NaOH (pH 7.5), 150 mM KCl, 5% (v/v) glycerol, and 1 mM TCEP. Protein was analyzed for purity by 10-20% SDS-Page (Figure S1) and then dialyzed overnight into the storage buffer containing 20 mM HEPES–NaOH (pH 7.5), 100 mM KCl, 5% (v/v) glycerol, and 1 mM TCEP. All proteins were finally concentrated, flash frozen in liquid nitrogen, and stored at −80°C. All associated plasmids and vectors can be found in Tables 2 and 3, respectively.

### Cryo-EM Preparation, Data Collection, and Data Processing

The type I-C Cascade was mixed with dsDNA target at a 1:2 molar ratio (complex:dsDNA). Target binding was facilitated by incubating the mixture at 30° C for 30 min. The CF-2/2 grids were first glow discharged for 60s and then a layer graphene oxide was added^33^. 3uL of protein was deposited on the grid and excess protein was blotted away after a 0.5s incubation time for 4s using filter paper at 4 °C in 100% humidity. The grid was then plunge frozen into liquid ethane using a Vitrobot Mark IV (Thermo Fisher). Frozen-hydrated samples of type I-C Cascade were directly visualized using a FEI Titan Krios microscope equipped with a Gatan K3 direct electron detector. Using the automated data-collection software LEGINON^36^, we acquired 5,399 movies at a magnification of x22,500, corresponding to a calibrated pixel size of 1.045 Å/pixel, a dosage of 15 e^-^/pixel/s, and a defocus range of −1.2 μm to −2.2 μm. Movies were collected on a Gatan K3 in 20 frames over an exposure time of 3 s (150 ms/frame), giving a total exposure of 45 e^-^/pixel. Data collected from the FEI Titan Krios was pre-processed in the real-time pre-processing tool WARP^37^. Motion correction, CTF-estimation and non-templated particle picking was performed in Warp^37^. After particle picking using WARP’s BoxNet, a neural network-based particle picker, 1.2 million particles and 5,399 micrographs were uploaded to cryosparc v2^38^.

For the Cascade-R-loop dataset, extracted particles were imported into CryoSPARC^38^ for 2D classification and 381,497 particles were selected for 3D classification *ab-initio* and subsequent hetero-refinement and non-uniform refinement. 778,052 particles were selected for 3D-classification to separate out dsDNA bound Cascade from ssDNA bound. Two distinct classes containing each a dsDNA and ssDNA bound state independently went through particle subtraction, CTF refine (G and L), a final non-uniform refinement. 174,004 particles went into a final reconstruction at 2.80-Å resolution of the type I-C Cascade bound to ssDNA. 96,964 particles went into a final reconstruction at 2.86-Å resolution of the type I-C Cascade bound to dsDNA, respectively. Both reconstructions were determined using the 0.143 gold standard Fourier Shell Correlation – calculated from two independent half-sets – criterion. The apo-model was docked in and used to assist *de-novo* building in Coot^39^, and refined in PHENIX^40^ and ISOLDE^41^. Alpha Fold^25^ was used to assist *de-novo* building of the N-term Cas8c in Coot^39^.

The type I-C Cascade was mixed independently with AcrIC4 and AcrIF2 at a 1:10 molar ratio (Cascade:Acr) and diluted to a concentration of 0.3mg/ml. Acr binding was facilitated by incubating the mixture at 30° C for 30 min. The CF-1.2/1.3 grids were first plasma cleaned for 30s and 2.5uL of sample was deposited on the grid. Excess protein was blotted away after a 0.5s incubation time for 6s with a force of 0 using filter paper at 4 °C in 100% humidity. The grid was then plunge frozen into liquid ethane using a Vitrobot Mark IV (Thermo Fisher). Frozen-hydrated samples of type I-C Cascade bound to AcrIC4 and AcrIF2 were directly visualized using a FEI Glacios microscope equipped with a Gatan K3 direct electron detector operating at 200kV. Images were taken at a pixel size of 0.94 Å/pixel using a Gatan K3 direct electron detector and a final dosage of 40.5 e^-^/Å^2^. Data collection was automated using SerialEM using a defocus range of -1.2 to -2.2 µm. 4,078 and 3,520 movies were collected from the Gatan K3 of type I-C Cascade bound to IF2 and IC4, respectively, and uploaded to cryoSPARC v2 Live^38^.

After motion and CTF correction, initial templates for template-based picking were generated using the apo-model in cryosSPARC v2 Live. Template-based particle picking of the type I-C Cascade bound to AcrIF2 and AcrIC4 resulted in 942,324 and 1.4 million particles, respectively. Processing for both datasets started three rounds of 2D classification, ab-initio, hetero-refinement, and non-uniform refinement. 160,716 and 128,780 particles of Cascade-AcrIF2 and Cascade-AcrIC4 datasets, respectively, were re-extracted and exposed to CTF refinement (G and L) and 3D-Classification using a mask over the Acrs to improve the quality of the map. A final round of non-uniform refinement and CTF refinement (G and L) resulted in a final model composed of 21,625 particles at a 3.0-Å resolution at for the type I-C Cascade-AcrIF2 complex, and a final model composed of 21,651 particles at a 3.1-Å resolution for type I-C Cascade-AcrIC4 complex. Both reconstructions were determined using the 0.143 gold standard Fourier Shell Correlation – calculated from two independent half-sets – criterion. The apo-model was docked in to both cryo-EM map and a previous structures of AcrIF2 was docked into the IF2-bound cryo-EM map. Alpha Fold^25^ was used to assist *de-novo* building of AcrIC4 in Coot^39^. Both final structures were refined in PHENIX^40^ and ISOLDE^41^.

To improve our original 3.1-Å resolution structure of the apo-model, we diluted the type I-C Cascade to 0.3mg/ml and followed the same cryo-EM preparation and data collection parameters as above (with Acrs). Template-based particle picking of the type I-C resulted in 3 million particles and processing followed the same protocol as listed above (with Acrs). Our previous apo type I-C Cascade model was docked into a new 2.80-Å map and used as a template to assist *de-novo* building. The final model was refined in PHENIX^40^ and ISOLDE^41^.

### Electrophoretic Mobility Shift Assays

Gel shift assays were performed in 1× binding buffer [50 mM Tris-HCl (pH 8.0), 150 mM NaCl, and 0.03% tween]. WT and mutant Cascades were diluted into 1× binding buffer to concentrations of 10nM, 20nM, 40nM, 80nM, 160nM, 320nM, 640nM, and1280nm. An assembled dsDNA duplex containing a 5’FAM-TS and complimentary NTS was added to a final concentration of 10 nM. Varying concentrations of Cascade were incubated with dsDNA at 37°C for 30 min and resolved at 4°C on 1% agarose containing 1× TBE. DNA was visualized by fluorescence imaging and images were quantified using ImageJ software. The fraction of DNA bound (amount of bound DNA divided by the sum of free and bound DNA) was plotted versus the concentration of type I-C Cascade and fit to standard one-site binding isotherm (all R-square values ≥ 0.98) using Prism (GraphPad). Reported KD values are the average of at least three independent experiments, and error bars represent the standard deviation.

### In-Vivo Interference Assays

The *in vivo* interference assay was adapted from the plasmid system from Dillard, et al., 2018^31^. All genes necessary for the formation and assembly of the *D. vulgaris* type I-C Cascade (Cas7c-Cas8c-Cas5c from addgene plasmid #81185)^16^ and associated cas3 (AAS94335.1) were cloned into a pBAD-based vector with ampicillin antibiotic resistance. Plasmid #81185 was used as associated type I-C Cascade crRNA. The 35 base pair target sequence with a 5’ TTC PAM site were cloned into a pCDF-Duet1 vector with streptomycin resistance. AcrIF2 (addgene and AcrIC4 were cloned into a pET28b vector with kanamycin resistance, respectively. LB agar plates were prepared with the following antibiotic concentrations: 50 µg/ml kanamycin, 100 µg/ml carbenicillin (or ampicillin), 50 µg/ml streptomycin, and 34 µg/ml chloramphenicol.

Vectors harboring the crRNA and target sequenced were co-transformed into BL21-AI cells, and were made electrocompetent with a series of glycerol washes. The remaining vectors were transformed using electroporation in order to obtain three E. Coli strains containing: crRNA+target+IC_cascade_cas3, crRNA+target+IC_cascade_cas3+AcrIF2, or crRNA+target+IC_cascade_cas3+AcrIC4. Single colonies of E. coli with crRNA+target+IC_cascade_cas3 were inoculated into 5ml of LB containing the appropriate antibiotics and grown overnight at 37° shaking at 225 rpm. The following day, the cells were centrifuged at room temperature at 3000 rpm for 10 minutes. The cells were decanted and resuspended with 5ml of LB with no antibiotics. A 1:100 dilution of the resuspended overnight cell cultures was inoculated into fresh LB with no antibiotics and grown to an OD 0.5 at 37° shaking. Cells were induced with a final concentration of 0.5% L-arabinose and grown for an additional 4 hours at 37°. Cell cultures were then serially 10-fold diluted and plated on LB agar plates containing chloramphenicol and carbenicillin, and chloramphenicol, carbenicillin, and streptomycin. Cleavage efficiency was calculated by the ratio of colony forming units (CFUs) on plates without streptomycin and with streptomycin. This was repeated for single colonies of E. coli harboring the crRNA+target_IC_cascade_cas3+AcrIF2 and crRNA+target+IC_cascade_cas3+AcrIC4. The induced cultures from these strains were serially 10-fold diluted and plated on LB agar plates containing chloramphenicol, carbenicillin and kanamycin, and chloramphenicol, carbenicillin, kanamycin, and streptomycin. Cleavage interference of the anti-crispr was calculated by the ratio of colony forming units (CFUs) on plates without streptomycin and with streptomycin. All associated plasmids and vectors can be found in Tables 2 and 3, respectively. Each sample was comprised of 15 biological replicates.

Welch’s t-test was used to calculate statistical significance of the cleavage efficiency of cell lines harboring each anti-crispr, respectively, compared to the cleavage efficiency of cell lines harboring IC-cascade and cas3 without the addition of an anti-crispr. The cleavage efficiency between E. coli cells without an anti-crispr and cells containing either acrIF2 or acrIC4 are statistically significant (Acr_IF2: P < 0.0001, Acr_C4: P < 0.0001)

## Acknowledgements

We thank Axel Brilot for expert cryo-EM assistance and members of the Taylor lab for helpful discussions. We especially thank Evan Schwartz and Isabel Strohkendl for insightful feedback on the manuscript. We also thank Ailong Ke and Ilya Finkelstein for the plasmids used to create our *in-vivo* interference assays. Data were collected at the Sauer Structural Biology Laboratory at the University of Texas at Austin. This work was supported by the National Institute of General Medical Sciences (NIGMS) of the National Institutes of Health (NIH) (R35GM138348) (to D.W.T). D.W.T is a CPRIT Scholar supported by the Cancer Prevention and Research Institute of Texas (RR160088).

## Author contributions

R.E.O. purified and reconstituted the type I-C Cascade complex. R.E.O purified all type I-C Cascade mutants and performed all EMSAs. D.R. and G.N.H. cloned AcrIC4, and R.E.O and J.W. purified AcrIF2 and AcrIC4. D.R. completed all in-vivo interference assays. R.E.O reconstituted type I-C complex bound to dsDNA, AcrIF2, and AcrIC4 and collected all cryo-EM data. R.E.O. and J.P.K.B. processed and interpreted all the cryo-EM data. R.E.O. built and refined the cryo-EM models. R.E.O. and D.W.T. wrote the manuscript with input from all authors. D.W.T. conceived the experiments, supervised the research, and secured funding for the project.

## Competing interests

The authors declare no competing interests.

## Supplementary Information

**Supplementary Figure 1.**
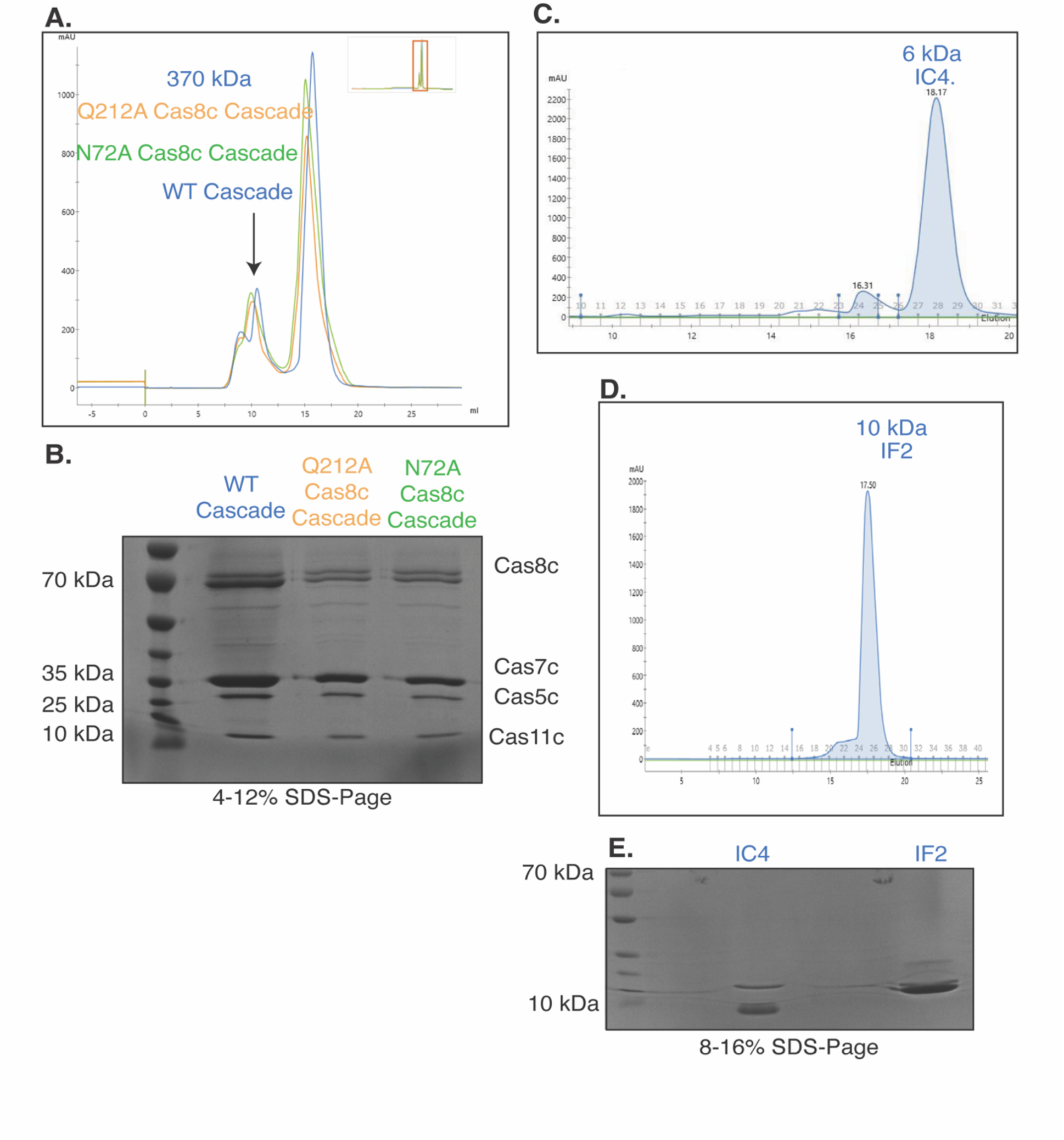
Purification of the type I-C Cascade Cas8c mutant Cascade, AcrIF2 and AcrIC4. a,b. Size-exclusion chromatograph and SDS-Page analysis of Cascade WT, Cas8c-Q212A Cascade, and Cas8c-N72A Cascade **d,e,f** Size-exclusion chromatograph and SDS-Page analysis of AcrIF2 and AcrIC4.

**Supplementary Figure 2.**
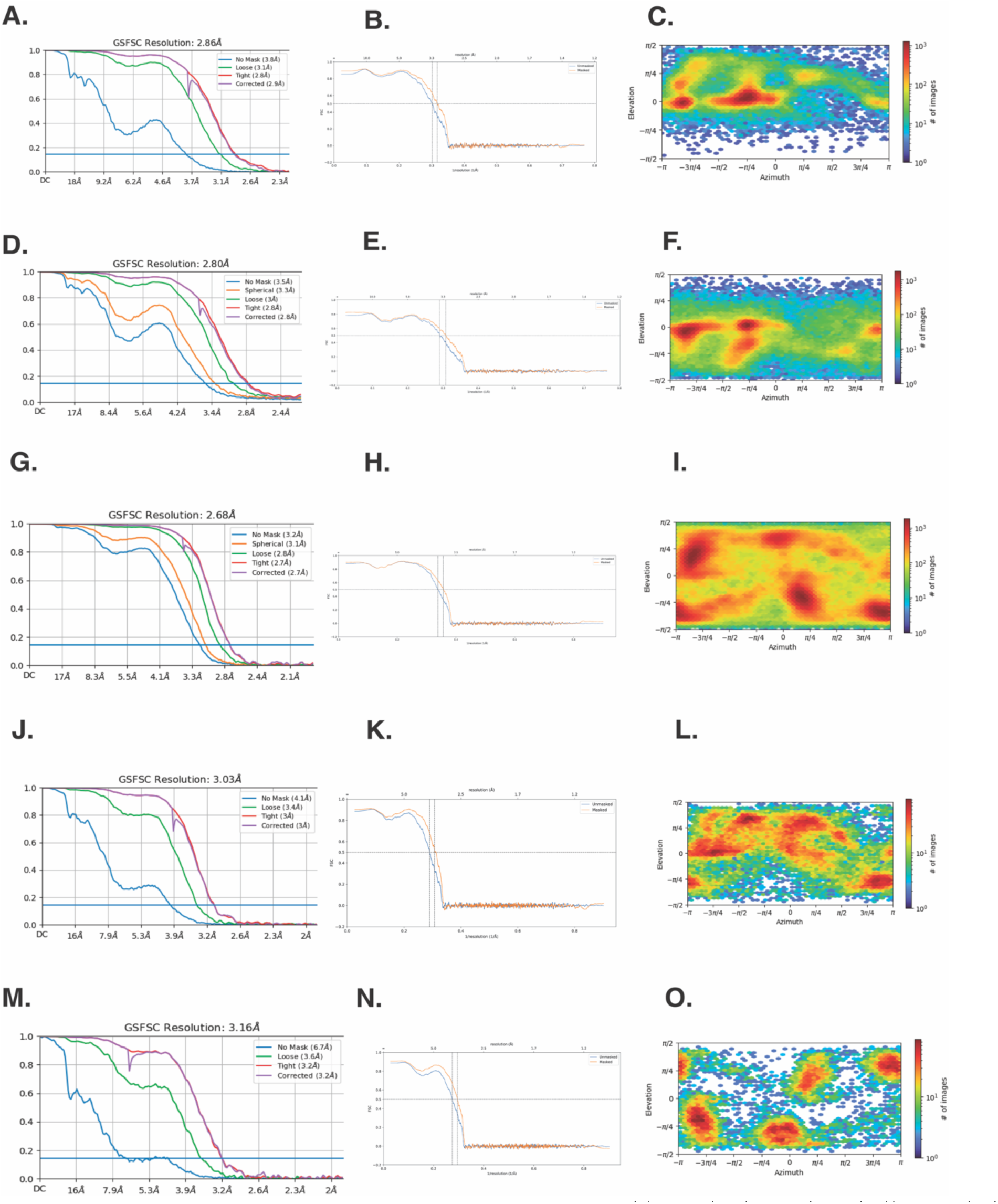
Cryo-EM data analysis. **a**, Gold standard Fourier Shell Correlation (FSC) curves for type I-C Cascade bound to dsDNA complex. **b,** Map-to-model FSC for type I-C Cascade bound to dsDNA complex **c**, Euler plot for type I-C Cascade bound to dsDNA complex **d**, Gold standard Fourier Shell Correlation (FSC) curves for type I-C Cascade bound to ssDNA complex. **e,** Map-to-model FSC for type I-C Cascade bound to ssDNA complex **f**, Euler plot for type I-C Cascade bound to ssDNA **g**, Gold standard Fourier Shell Correlation (FSC) curves for unbound type I-C Cascade. **h,** Map-to-model FSC for type I-C Cascade bound to unbound type I-C Cascade **i**, Euler plot for unbound type I-C Cascade **j**, Gold standard Fourier Shell Correlation (FSC) curves for type I-C Cascade bound to AcrIF2. **k,** Map-to-model FSC for type I-C Cascade for type I-C Cascade bound to AcrIF2. **l**, Euler plot for type I-C Cascade bound to AcrIF2. **m**, Gold standard Fourier Shell Correlation (FSC) curves for type I-C Cascade bound to AcrIC4. **n,** Map-to-model FSC for type I-C Cascade for type I-C Cascade bound to AcrIC4. **o**, Euler plot for type I-C Cascade bound to AcrIC4.

**Supplementary Figure 3.**
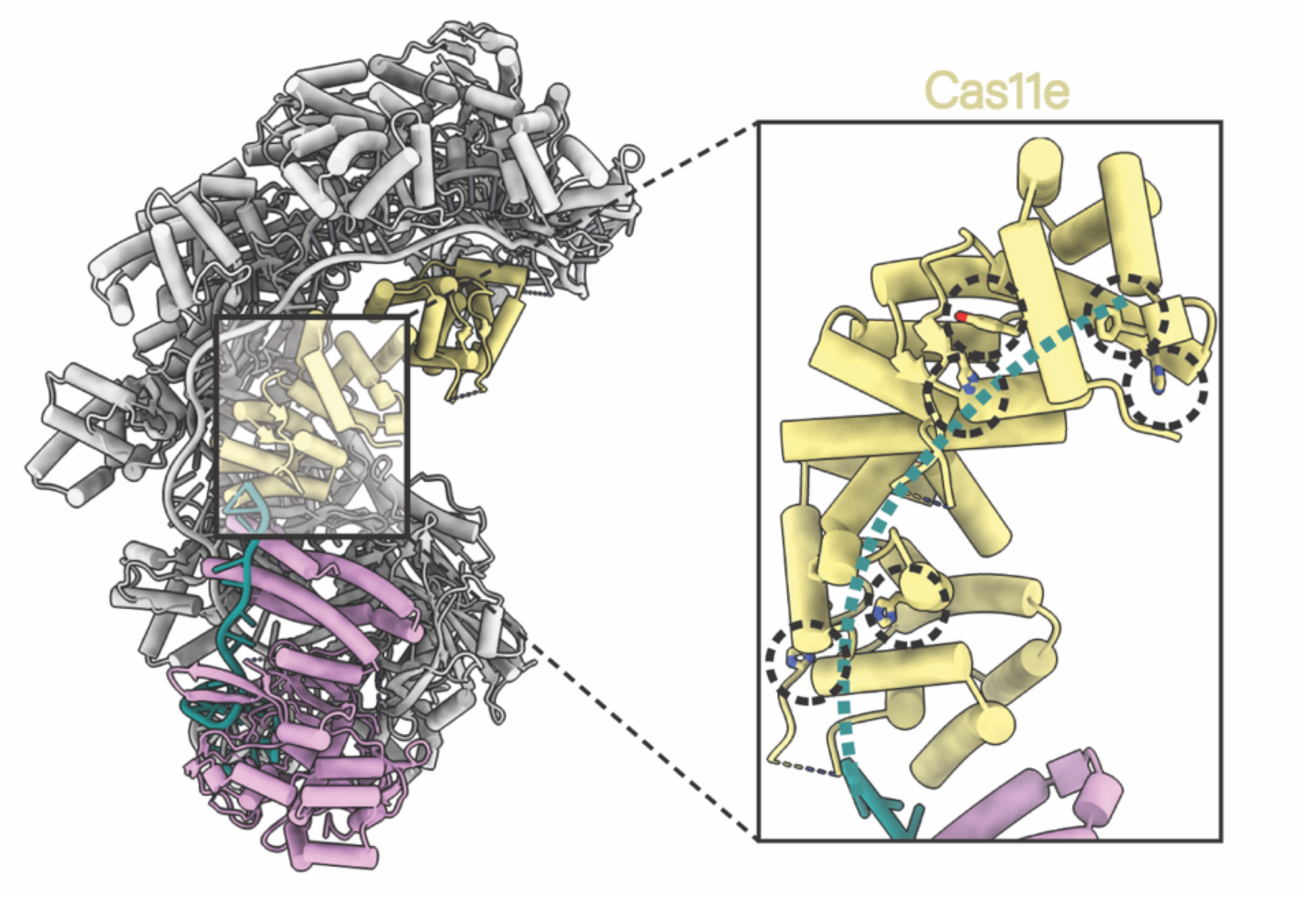
Multiple aromatic residues similarly positioned along the putative NTS path (green) of type I-E Cascade.

**Supplementary Figure 4.**
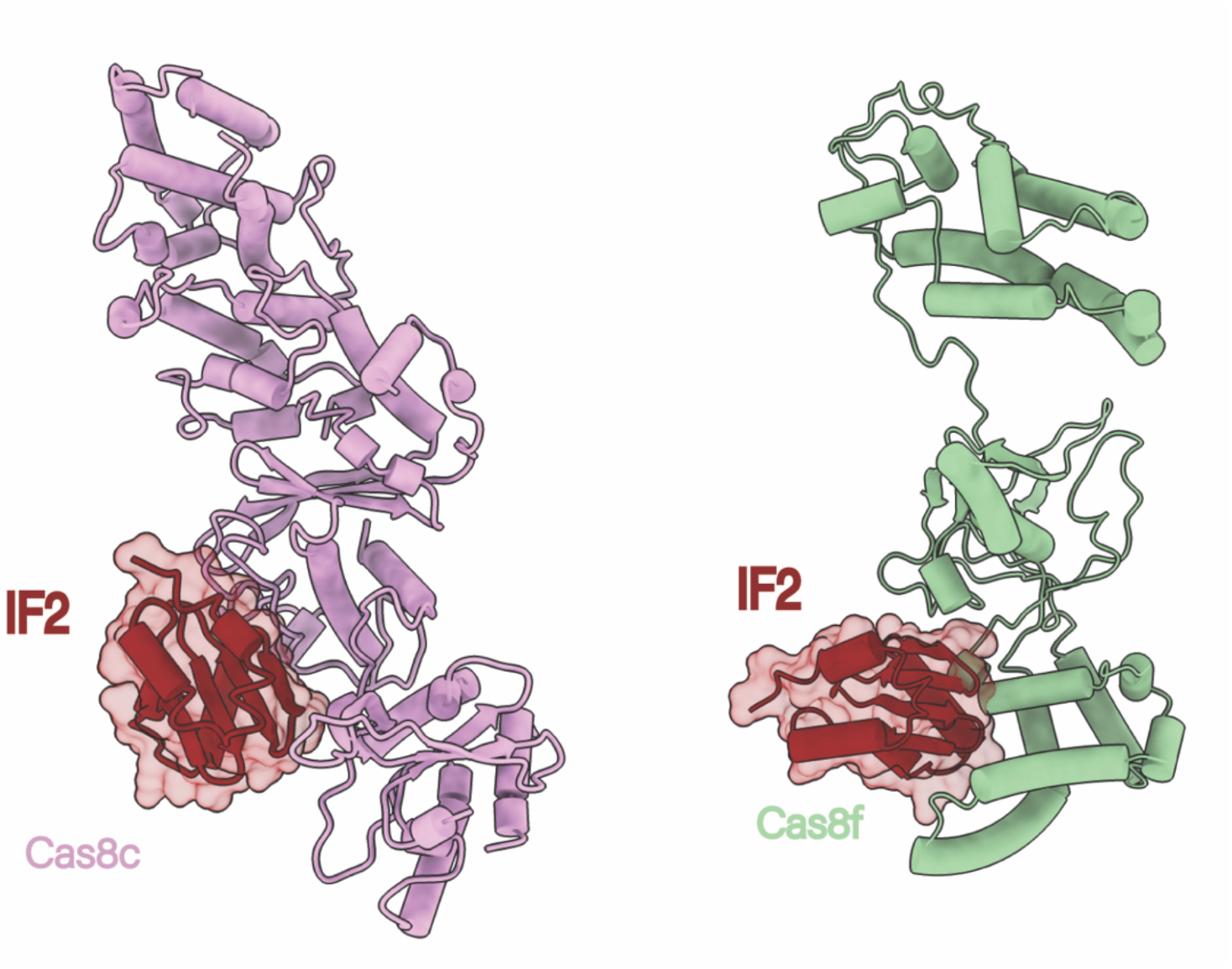
IF2 inhibits both Cas8c and Cas8f through the same interface but at different orientations.

**Supplementary Table 1:** Model Statistics for the type I-C Cascade structures.

|  | Type I-C<br>apo model<br>unbound | Type I-C<br>Cascade<br>bound to<br>dsDNA | Type I-C<br>Cascade<br>bound to<br>ssDNA | Type I-C<br>Cascade<br>bound to<br>AcrIF2 | Type I-C<br>Cascade<br>bound to<br>AcrIC4 |
| --- | --- | --- | --- | --- | --- |
| **Data collection and processing** |  |  |  |  |  |
| Voltage (kV) | 200 | 300 | 300 | 200 | 200 |
| Electron exposure (e-/Å<sup>2</sup>) | 40.5 | 80 | 80 | 40.5 | 40.5 |
| Defocus range (μm) | 1.2 to 2.2 | 1.2 to 2.2 | 1.2 to 2.2 | 1.2 to 2.2 | 1.2 to 2.2 |
| Pixel size (Å) | 0.94 | 1.047 | 1.047 | 0.94 | 0.94 |
| Symmetry imposed | C1 | C1 | C1 | C1 | C1 |
| Initial particle images (no.) | 2 million | 1.2 million | 1.2 million | 160,716 | 128,780 |
| Final particle images (no.) | 73,220 | 96,964 | 174,004 | 21,625 | 21,651 |
| Map resolution (Å) | 2.84 | 2.86 | 2.80 | 3.1 | 3.16 |
| FSC threshold | 0.143 | 0.143 | 0.143 | 0.143 | 0.143 |
| Map resolution range (Å) | N/A | N/A | N/A | N/A | N/A |
| **Refinement** |  |  |  |  |  |
| Initial pdb model used | 7KHA | 7KHA | 7KHA | 7KHA<br>5UZ9 | 7KHA |
| Model resolution (Å) | 2.8 | 3.2 | 3.2 | 3.3 | 3.4 |
| FSC threshold | 0.5 | 0.5 | 0.5 | 0.5 | 0.5 |
| Model resolution range (Å) | N/A | N/A | N/A | N/A | N/A |
| Map sharpening *B* factor (Å<sup>2</sup>) | 89.3 | 120.3 | 127.1 | 60.0 | 47.5 |
| Model composition |  |  |  |  |  |
| Non-hydrogen atoms | N/A | N/A | N/A | N/A | N/A |
| Protein residues | 2843 | 2975 | 23796 | 3063 | 2900 |
| Ligands | 48 | 133 | 68 | 48 | 46 |
| *B* factors (Å<sup>2</sup>) |  |  |  |  |  |
| Protein | 60.78/283.4<br>5/117.35 | 45.35/475.79/<br>133.24 | 73.81/724.86/<br>195.04 | 50.90/296.18/<br>119.13 | 61.57/335.78/<br>130.94 |
| Nucleotide | 70.28/197.1<br>6/108.60 | 64.05/501.15/<br>168.17 | 95.29/343.01/<br>162.77 | 60.54/208.99/<br>106.80 | 84.37/205.02/<br>121.24 |
| R.m.s. deviations |  |  |  |  |  |
| Bond lengths (Å) | .0004 | .004 | .003 | 0.003 | 0.003 |
| Bond angles (°) | .614 | .673 | .682 | 0.633 | 0.646 |
| **Validation** |  |  |  |  |  |
| MolProbity score | 1.52 | 1.56 | 1.76 | 1.53 | 1.62 |
| Clashscore | 6.02 | 5.65 | 11.00 | 7.57 | 8.52 |
| Poor rotamers (%) | 1.22 | 0.72 | 0.21 | 0.27 | 0.07 |
| Ramachandran plot |  |  |  |  |  |
| Favored (%) | 97.35 | 96.27 | 96.80 | 97.44 | 97.09 |
| Allowed (%) | 2.65 | 3.73 | 3.20 | 2.56 | 2.84 |
| Disallowed (%) | 0.00 | 1.00 | 0.00 | 0.00 | 0.07 |

**Supplementary Table 2:** Plasmids used in this study.

| **Plasmid** | **Description** | **Reference** |
| --- | --- | --- |
| #81185 | *D. vulgaris* type I-C Cascade for cryo-EM studies | Addgene |
| #81186 | *D. vulgaris* type I-C crRNA for co-expression with Cascade | Addgene |
| #89234 | AcrIF2 for purification and cryo-EM studies | Addgene |
| pIF199 | Backbone used for AcrIF2 and AcrIC4 for *in vivo* interference assay | Dillard et al |
| pIF412 | Backbone used for dsDNA target for *in vivo* interference assay | Dillard et al |
| pIF385 | Backbone used for Cascade-Cas3 for *in vivo* interference | Dillard et al |
| #48313 | Backbone vector used for AcrIC4 for *in vivo* interference | Addgene |
| **pRO01** | Type I-C Cascade Cas8c mutant Q212A for purification and EMSA | This study |
| pRO02 | Type I-C Cascade Cas8c mutant N72A for purification and EMSA | This study |
| pRO03 | AcrIC4 for purification and cryo-EM studies | This study |
| pIS010 | pACYC vector with IC crRNA array *in vivo* interference | This study |
| pIS011 | pCDF-Duet1 vector with target sequence and 5' PAM for *in vivo* interference | This study |
| pDR005 | pBAD vector with IC cascade and cas3 for *in vivo* interference | This study |
| pDR007 | pET28b-Duet1 vector with Acr IF2 for *in vivo* interference | This study |
| pDR008 | pET28b-Duet1 vector with Acr-IC4 for *in vivo* interference | This study |

**Supplementary Table 3:**
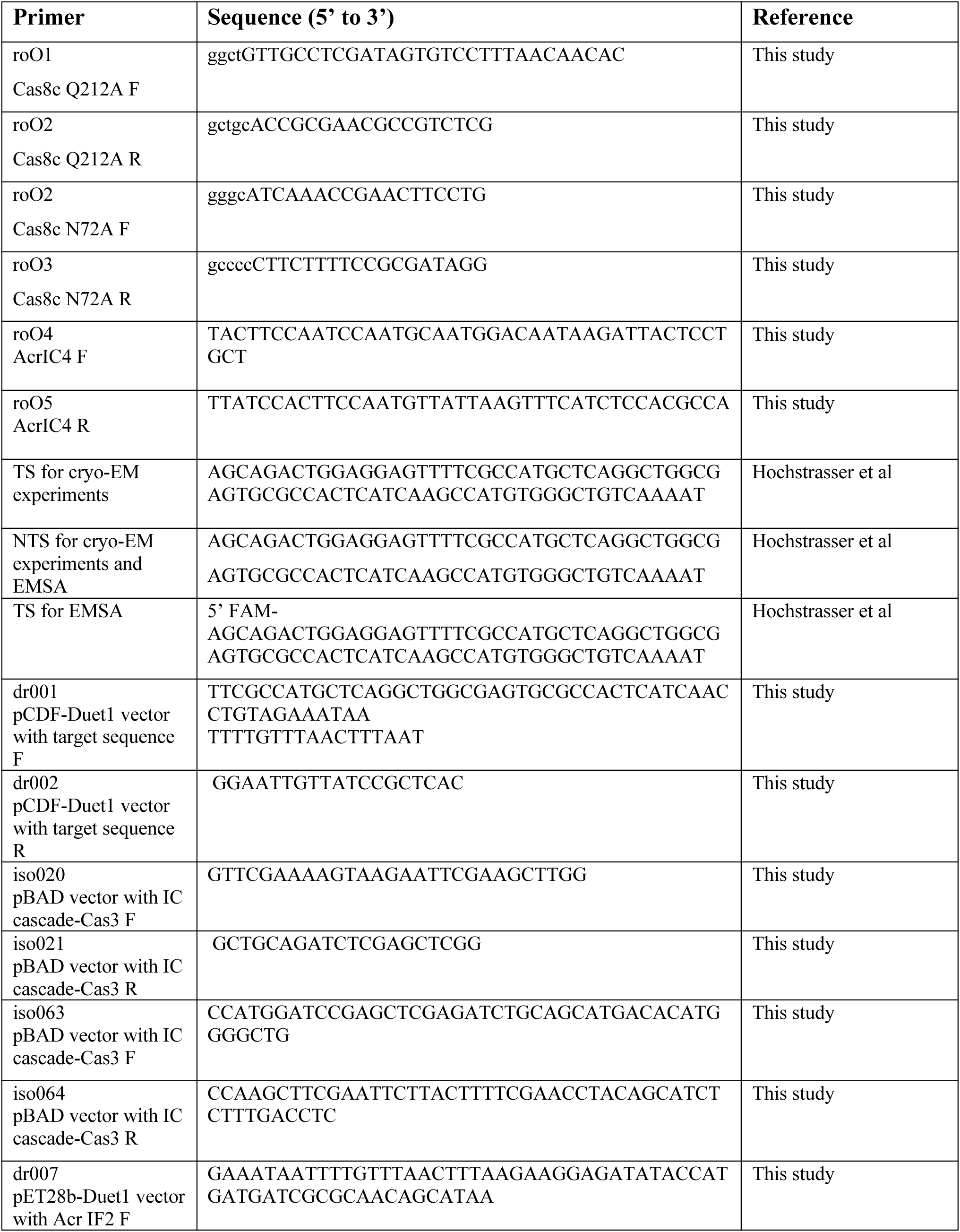
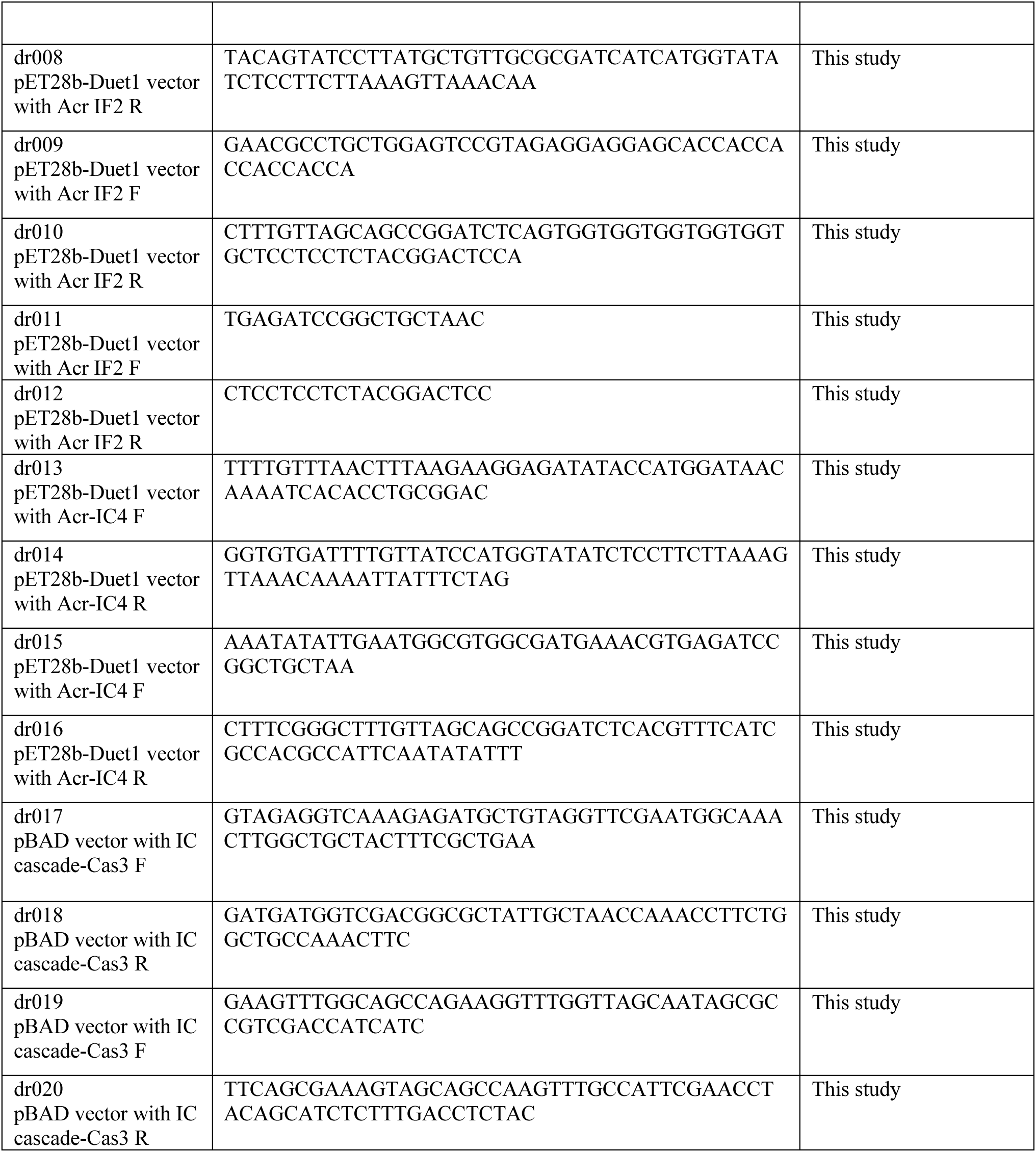
Oligonucleotides used in this study.

## Notes

### Competing Interest Statement

The authors have declared no competing interest.

